# ALOHA: AI-guided tool for the quantification of venom-induced haemorrhage in mice

**DOI:** 10.1101/2022.08.04.502754

**Authors:** Timothy P. Jenkins, William Michael Laprade, Andrés Sánchez, Tulika Tulika, Carol O’Brien, Christoffer V. Sørensen, Trenton K. Stewart, Thomas Fryer, Andreas H. Laustsen, José María Gutiérrez

## Abstract

Venom-induced haemorrhage constitutes a severe pathology in snakebite envenomings, especially those inflicted by viperid species. In order to both explore venom compositions accurately, and evaluate the efficacy of viperid antivenoms for the neutralisation of haemorrhagic activity it is essential to have available a precise, quantitative tool for empirically determining venom-induced haemorrhage. Thus, we have built on our prior approach and developed a new AI-guided tool (ALOHA) for the quantification of venom-induced haemorrhage in mice. Using a smartphone, it takes less than a minute to take a photo, upload the image, and receive accurate information on the magnitude of a venom-induced haemorrhagic lesion in mice. This substantially decreases analysis time, reduces human error, and does not require expert haemorrhage analysis skills. Furthermore, its open access web-based graphical user interface makes it easy to use and implement in laboratories across the globe. Together, this will reduce the resources required to preclinically assess and control the quality of antivenoms, whilst also expediting the profiling of hemorrhagic activity in venoms for the wider toxinology community.

## 1. Introduction

Snakebite envenoming is a major public health problem, especially in the developing world [1]. Indeed, it is responsible for substantial morbidity and mortality, particularly in the impoverished areas of sub-Saharan Africa, South to Southeast Asia, Papua New Guinea, and Latin America [1–4]. Whilst accurate estimates are difficult to make, it is believed that between 1.8–2.7 million people worldwide are envenomed each year, resulting in 80,000 to 140,000 deaths and 400,000 survivors left with permanent sequelae [5–7].

The severity of a given envenoming is determined by several factors, such as the amount of venom injected, the anatomical location of the bite, and the physiological status of the victim [8]. In addition, there is a great variability in the composition of the venoms and the predominant toxins present in different venoms, not only between genera, but also within a single species [9]. Consequently, the clinical manifestations and pathophysiological effects of envenomings can vary greatly depending on the offending snake species [10]. One such effect is haemorrhage. Indeed, envenomings by many snakes, predominantly by species belonging to the family Viperidae, but also some from the family Colubridae (*sensu lato*), induce local and systemic haemorrhage, further causing local tissue damage and cardiovascular disturbances [8,11,12]. Specifically, blood vessel damage leads to extravasation, which contributes to local tissue damage and poor muscle regeneration. In addition, massive systemic haemorrhage contributes to hemodynamic disturbances and cardiovascular shock [8,13]. Consequently, the underlying mechanisms by which snake venoms induce haemorrhage, the characterisation of hemorrhagic toxins, and the clinical manifestations of envenomings have, for a long time, presented a key area of fundamental, but also translational research within the field of Toxinology [11]. Venom-induced haemorrhage is mainly the consequence of the damage induced by snake venom metalloproteinases (SVMPs) on the microvasculature, due to the enzymatic degradation of key structural components in the basement membrane of capillary vessels [12]. The haemorrhagic activity of venoms is further potentiated by the action of venom toxins that affect haemostasis, which induce consumption coagulopathy, thrombocytopenia, and platelet hypoaggregation [1,14].

The only specific treatment currently available for snakebite envenomings is the intravenous administration of animal-derived antivenom [8,15]. Importantly, each batch of antivenom that is produced needs to undergo rigorous quality control; this includes the assessment of the ability of antivenoms to neutralise the lethal effect of venoms in mice [16,17]. In addition, and owing to the complex pathophysiology of snakebite envenomings, other relevant effects, such as the antivenom’s neutralising potential of venom-induced haemorrhage, are also part of the preclinical assessment of antivenom efficacy [16]. The most widely used method for analysing haemorrhage is the skin test originally developed in rabbits [18], and later on adapted for use in rats [19] and mice [20]. In the adaptation of this method for mice, a range of different venom concentrations are injected intradermally in the abdominal region. After a predefined time interval, mice are euthanised and carefully dissected in order to allow the assessment of the inner surface of the skin. Originally, this was followed solely by a rough manual measurement of the area of the haemorrhagic lesion. Since this method did not take into account the intensity of the lesion, a computationally assisted update to this method was reported in 2017, in which an image of the lesion is taken, and both the size and the intensity are measured accurately, thus providing a more systematic and quantitative evaluation of extravasation [21]. The study also introduced a new unit for the assessment of the severity of a given venom-induced hemorrhagic lesion, i.e. the haemorrhagic unit (HaU) [21]. Whilst this method presented an improvement in accuracy and speed of the quantification of snake venom-induced haemorrhagic activity, it still required manual identification of the lesions and specialised equipment. It thus remained subject to human error and was not optimally accessible to researchers across the globe.

Thus, we present a new and more accessible artificial intelligence (**A**I)-guided too**l** for the automatic assessment **o**f venom-induced **ha**emorrhage, ALOHA **(https://github.com/laprade117/ALOHA)**. We trained a machine learning algorithm to automatically identify haemorrhagic lesions, adjust for lighting biases, scale the image, extract lesion area and intensity, and calculate the HaUs. Finally, we evaluate the performance of this algorithm and discuss its utility in relation to rapid, robust, and semi-automated assessment of snake venom-induced haemorrhage.

## 2. Methods

### 2.1 Snake venom

The venom of *Bothrops asper* was used in this study since its haemorrhagic activity has been widely studied. Venom of *B. asper* (batch number 03–06 Bap P) was obtained from adult specimens captured in the Pacific region of Costa Rica and maintained in captivity at the Serpentarium of Instituto Clodomiro Picado, Universidad de Costa Rica, San José, Costa Rica. Samples of venom correspond to pools obtained from many adult specimens and were stabilised by lyophilisation and stored at −20 °C. Solutions of venoms in 0.12 M NaCl, 0.04 M phosphate, pH 7.2 buffer (PBS) were prepared immediately before use.

### 2.2 Haemorrhagic activity

Haemorrhagic activity was assessed following the method described by Jenkins et al. (2017) with some modifications (c.f. below). Briefly, groups of four mice of both sexes (18–20 g; CD-1 strain) were injected intradermally with different amounts of *B. asper* venom (1, 2, 4, 8, 16 μg) dissolved in 100 μL PBS. Two hours after injection, mice were sacrificed by CO_2_ inhalation, and their skin was dissected. Mice were first placed on a standardised A4 printout template sheet to measure the haemorrhagic lesion on the inner surface of the skin, using the method described below; then the same mice were also placed on the table without the template printout sheet, with pictures being taken for both approaches. All experiments involving the use of mice were approved by the Institutional Committee for the Care and Use of Laboratory Animals (CICUA) of the University of Costa Rica (approval number CICUA 82-08).

### 2.3. Printout sheet

To allow for a standardised analysis of the haemorrhagic lesions, as well as to facilitate the image analysis algorithms, we prepared an A4 printout sheet which the mice were placed on. This sheet outlines where to place the mice and includes different lines and boxes of defined lengths that allow for the scaling of the image (Fig.1). We also used a cut out mask to be placed on the mice to facilitate lesion identification (Fig.1). Printable versions of these two components can be found in supplementary Figure 1 and 2.

**Figure 1.**
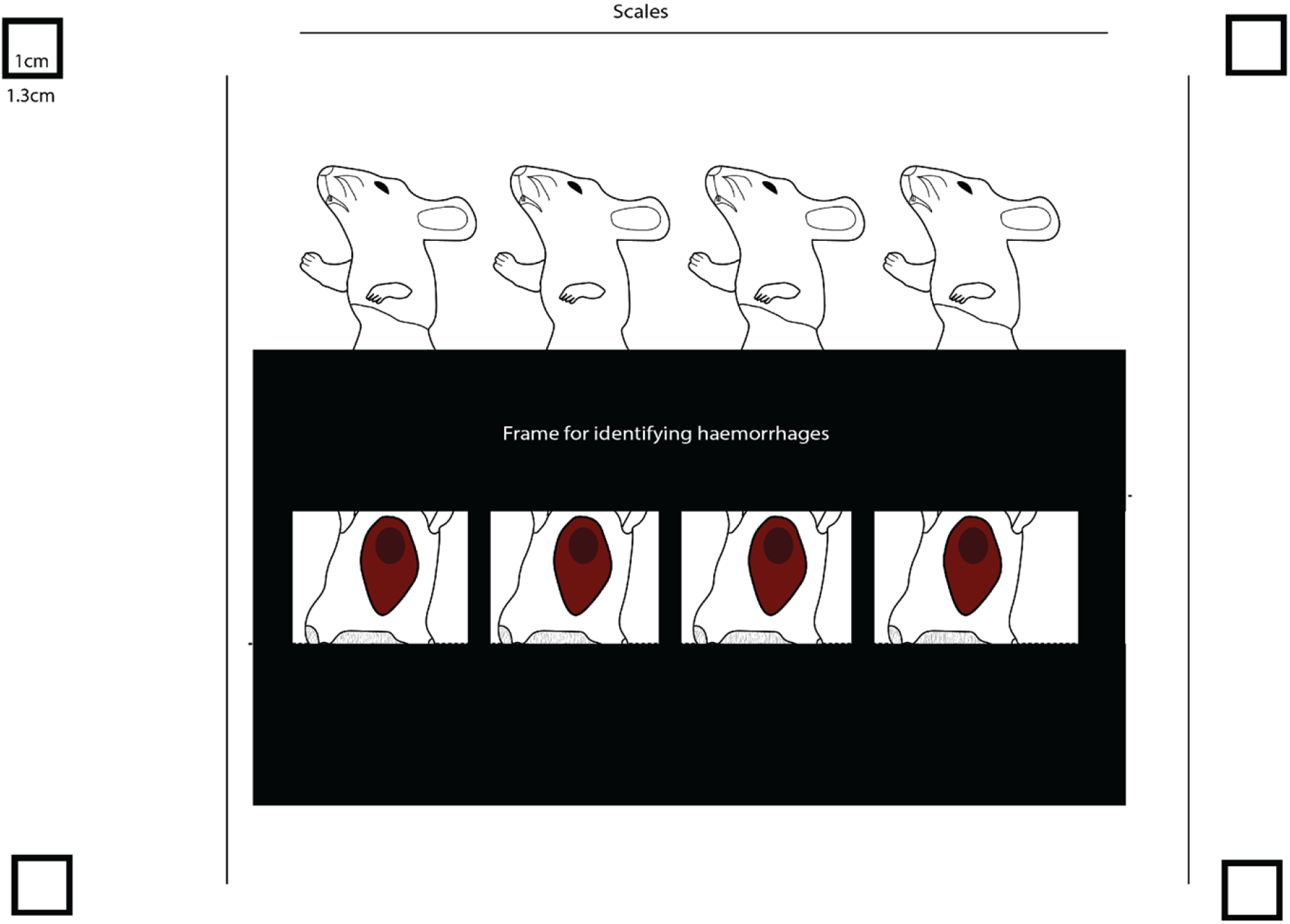
Explanation of the printable template upon which the mice should be placed. The multiple scales across the page allow for the automatic scaling by the tool and take into account pictures obtained from different angles. The black frame acts as a cut out mask to facilitate the automatic identification of haemorrhagic lesions.

### 2.4. Description of machine learning guided approach of quantifying haemorrhagic activity

We trained a machine learning algorithm to automatically identify haemorrhagic lesions, adjust for lighting biases, scale the image, extract haemorrhagic lesion area and intensity, and calculate the HaUs. This was then implemented in an accessible fashion via a graphical user interface (GUI) as the ALOHA tool (Fig. 2).

**Figure 2.**
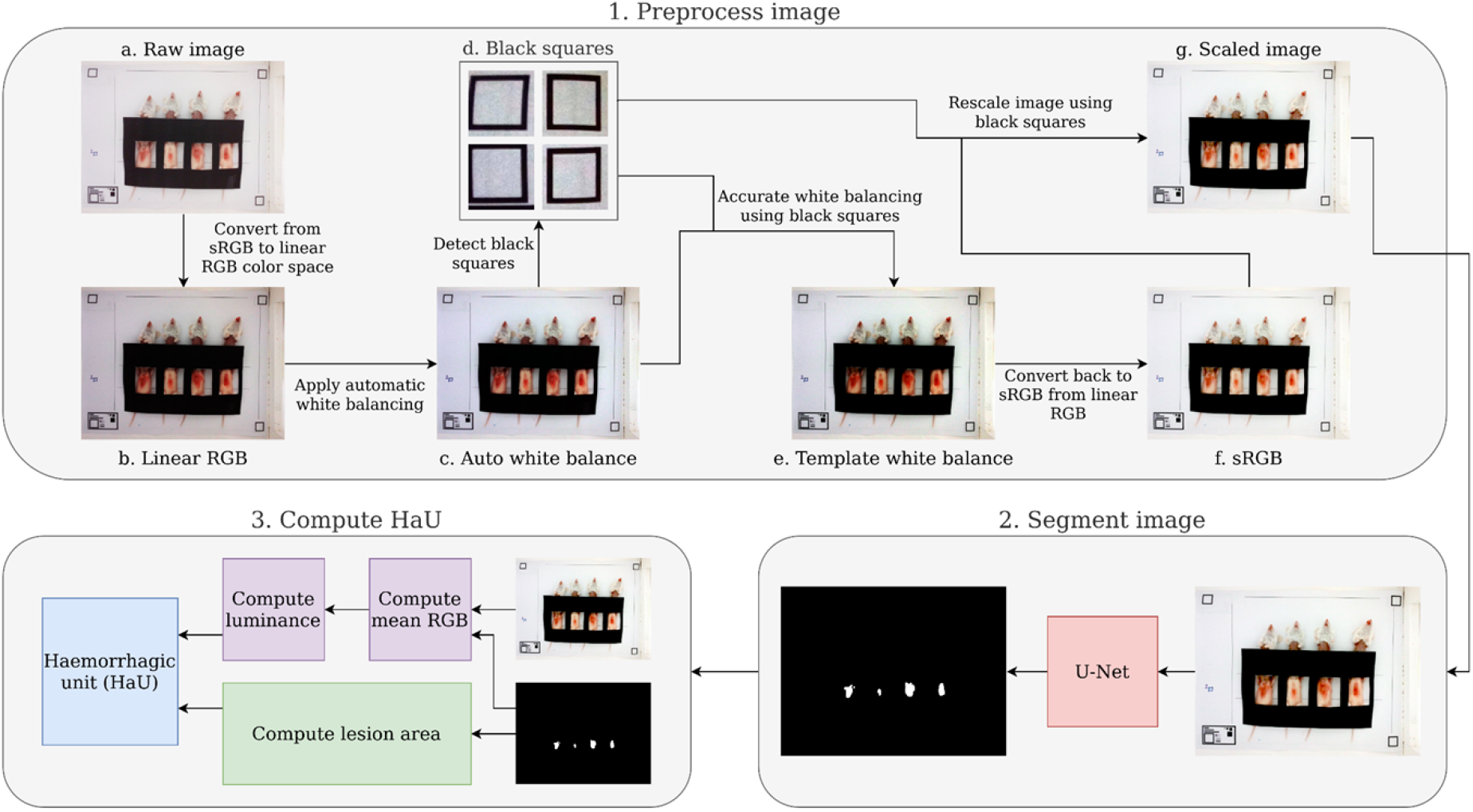
Overview of the workflow for ALOHA. First, the raw image is imported and converted from sRGB to linear RGB. Thereafter, the image is white balanced and subsequently further white balanced using the colour of the paper detected via the scaling squares. In parallel, the image is rescaled using the same squares. This processed image is then used for segmentation and automatic identification of the haemorrhagic lesions. Together, this information is used to compute the lesion area and luminance, which is then combined into a HaU score to assess the overall hemorrhagic lesion.

#### 2.4.1. Conversion to linear RGB and white balancing

In order to create reproducible results across images, it is necessary to white balance the images prior to computing the HaUs. First, the images were converted to a linear RGB colour space as in Jenkins et al. 2017 [21], then auto white balancing, based on the method used in the GNU Image Manipulation Program (https://www.gimp.org), was applied. For images that use our template, we applied a second white balancing step for higher accuracy using information extracted from the template image. Firstly, we detected the black boxes in the four corners of the image using a template matching algorithm. We then computed a white point based upon the mean colour of the pixels located within each of the four boxes and a black point based upon the mean colour of the pixels in the black borders of these boxes. The image is then white balanced via the following formula:

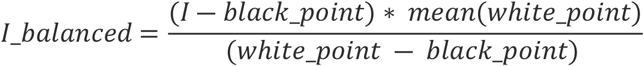

This formula does require the assumption that the paper which the template has been printed on is some value of grey, (i.e. the RGB values are all identical).

#### 2.4.2. Segmentation

To identify and segment the haemorrhagic lesions in images, a deep learning method based on the widely-used U-Net architecture was applied [22]. We also included a few modifications based upon more recent findings. Namely, we replaced the deconvolution layers in the expanding path with bilinear upsampling followed by a 2×2 convolution and included batch normalisation layers [23,24]. The resulting architecture contains approximately 31 million trainable parameters.

Our dataset consisted of 29 training images taken via smartphone. Each image contains between 1 and 32 mice displaying varying levels of haemorrhagic damage for a total of 217 mice. To limit annotator bias, each image was annotated by two different annotators, resulting in two masks per image. For evaluating performance, we set aside 20% of the images at random as a validation test set. Thereafter, a 5-fold cross-validation was performed on the remaining 23 images, evaluating model performance on the validation test set for each fold. This was repeated 5 times to avoid test-set bias.

At the time of training, the images were split into samples of size 256 x 256 pixels and fed into the model in batches of 32 samples. Batches were created such that each sample had a 75% chance of having a masked section of haemorrhagic tissue according to at least one annotator. The masks used for training were sampled from the set of annotators at random. Data augmentations included flips, rotations, noise, blurring, sharpening, distortions, brightness, contrast, hue, and saturation adjustments. Augmentations were selected to simulate the possible variation in both the lighting environment as well as the smartphone camera’s built-in post-processing implementations.

The models were trained using the Adam optimizer with a learning rate of 0.0001 for 100 epochs. We used a loss function based upon a combination of the Mathews correlation coefficient (MCC) and cross-entropy [25]. We report the average F1 (Dice), MCC, and accuracy for each model as computed on the test set. F1/Dice is the harmonic mean of precision and recall and ranges between 0 and 1, where 1 is a perfect score. MCC ranges from −1 to 1, with 1 being a perfect score.

#### 2.4.3. Scaling

It is essential for computing an accurate HaU that the scale of the images is determined. With the template, this can be done automatically by first detecting the black squares (via template matching) in the corners and then determining the number of white pixels inside the black square via thresholding. The inner white region is a 10 mm x 10 mm square, so the pixel resolution can be computed with the formula below.

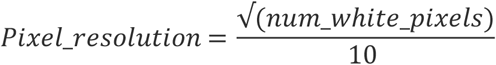

For improved accuracy, we then averaged the results across the four black squares.

For images without the template, the scale was determined manually by using FIJI’s measure tool on a known distance in the image [26]. Once the image scale was determined, we resized all images to ensure a resolution of 5 pixels per mm, which also allows for rapid computation.

#### 2.4.4 Calculation of HaU and minimum haemorrhagic dose

Haemorrhagic units were calculated as described in Jenkins *et al.* 2017 [21]. Briefly, the RGB values and area of a given lesion were extracted and colour/scale adjusted. Thereafter, the luminance (i.e., intensity) was calculated, combined with the area of the lesion, and expressed as HaUs. Thereafter, the minimum haemorrhagic dose (MHD) of *B. asper* venom was calculated. This calculation was carried out using linear regression on the means of the values from table 4. From the resulting function, we calculated the venom dose needed for a 50 HaU signal by replacing the Y with 50 and calculating X. The software used was GraphPad Prism version 9.2.0.

#### 2.4.5 Implementation in GUI

Using Streamlit (https://github.com/streamlit/streamlit) and localtunnel (https://github.com/localtunnel/localtunnel) with Google Colab, a simple web-based application to automatically analyse images was developed **(https://github.com/laprade117/ALOHA)**. A web-based application seems to be the most efficient way to quickly analyse data while working in the lab. Users can take a photo with a smartphone and upload it to the web-based tool (accessible via a smartphone browser) for an immediate result (supplementary Figure 3).

## 3. Results

In this study, we primarily present the results for the six images that used the template to evaluate the performance of our fully automated method, ALOHA. We used, as a model, the haemorrhagic lesions induced by the venom of *B. asper* on mice.

### 3.1 White balancing

To address the potential impact of lighting differences, the tool automatically performs white balancing. The white balancing works as expected and produces comparable results across images (Fig. 3).

**Figure 3.**
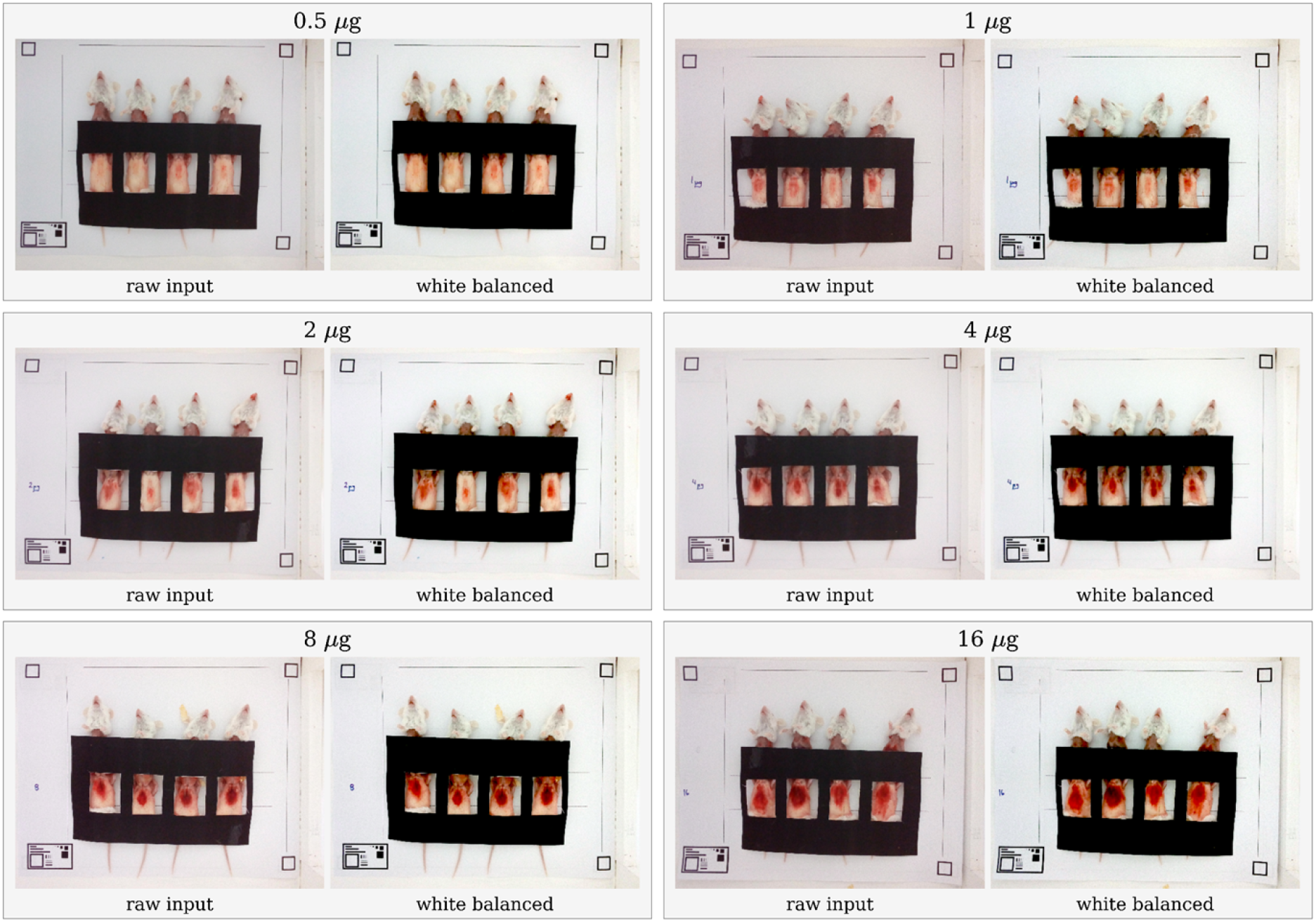
An overview of ALOHA’s automatic white balancing output across the different test images.

### 3.2 Scaling

Using the scaling method outlined in 2.4.3, the tool automatically detects the image scales and reports them (Table 1).

**Table 1.**
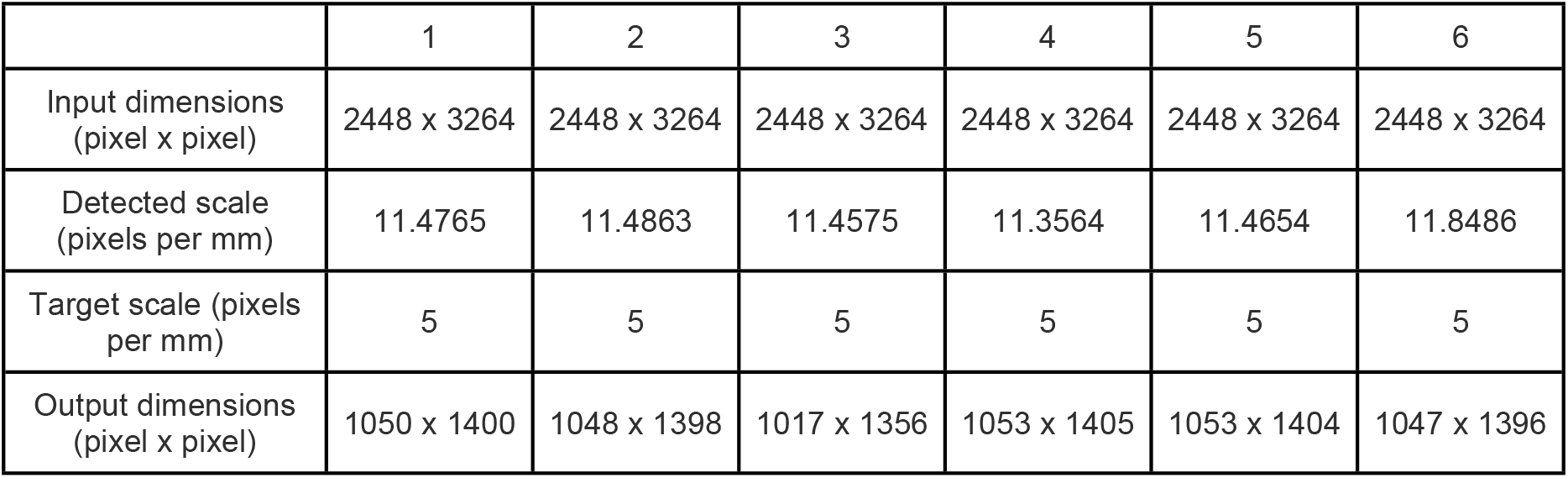
Table outlining the input dimensions, detected scale, target scale, and output dimensions across six test images.

|  | 1 | 2 | 3 | 4 | 5 | 6 |
| --- | --- | --- | --- | --- | --- | --- |
| Input dimensions (pixel x pixel) | 2448 x 3264 | 2448 x 3264 | 2448 x 3264 | 2448 x 3264 | 2448 x 3264 | 2448 x 3264 |
| Detected scale (pixels per mm) | 11.4765 | 11.4863 | 11.4575 | 11.3564 | 11.4654 | 11.8486 |
| Target scale (pixels per mm) | 5 | 5 | 5 | 5 | 5 | 5 |
| Output dimensions (pixel x pixel) | 1050 x 1400 | 1048 x 1398 | 1017 x 1356 | 1053 x 1405 | 1053 x 1404 | 1047 x 1396 |

### 3.3 Segmentation

To automatically identify lesion areas, the tool uses a machine learning guided segmentation approach. Overall, an average MCC score of 0.8612 and an average F1 (Dice) score of 0.9064 was achieved, and we were able to predict 99.84% of the pixels correctly across 25 runs (Fig. 4).

**Figure 4.**
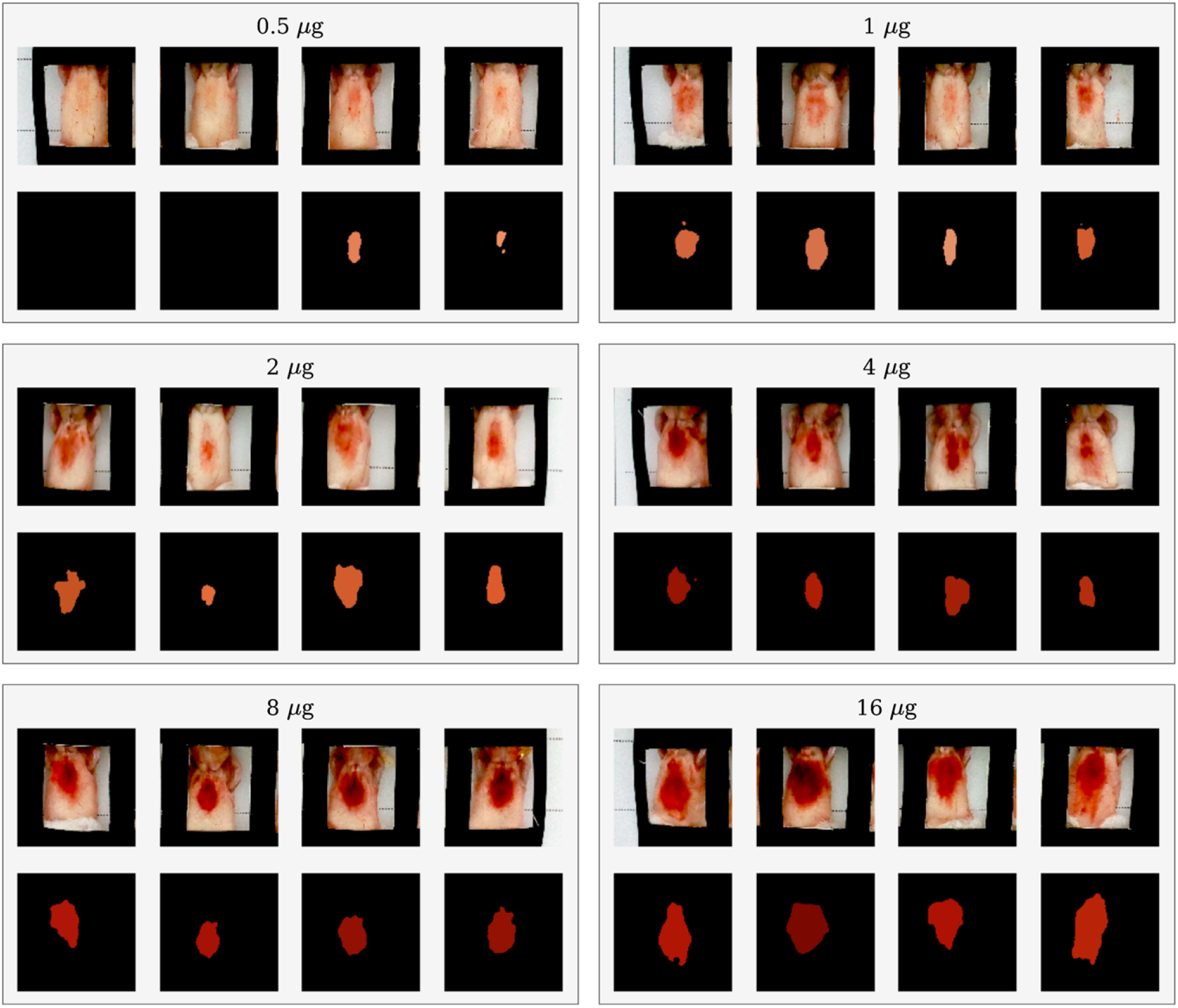
Segmentation of all haemorrhagic lesions across the test images. The segmentations are coloured using the average RGB values detected within the lesion area and the amount of *Bothrops asper* venom injected into the mice indicated above each batch of images.

### 3.4 Haemorrhagic Units

To assess the severity of each lesion, the tool automatically computes the real-world area, luminance, and HaU for each mouse in all of the test images (Tables 2,3,4).

**Table 2.** Individual and average (across one image) lesion sizes for all of the six test images. The amount of *Bothrops asper* venom injected into the mice is indicated next to each batch of images.

| Area (mm<sup>2</sup>) |  |  |  |  |  |  |
| --- | --- | --- | --- | --- | --- | --- |
| Venom (µg) | Mouse 1 | Mouse 2 | Mouse 3 | Mouse 4 | Average | Std. Dev. |
| 0.5 | 0.00 | 0.00 | 63.08 | 23.16 | 21.56 | 25.77 |
| 1 | 98.80 | 132.68 | 68.80 | 86.84 | 96.78 | 23.32 |
| 2 | 138.88 | 39.36 | 169.40 | 101.32 | 112.24 | 48.50 |
| 4 | 111.84 | 90.80 | 139.04 | 72.60 | 103.57 | 24.74 |
| 8 | 174.92 | 115.96 | 156.48 | 179.28 | 156.66 | 25.01 |
| 16 | 248.32 | 286.84 | 221.00 | 320.32 | 269.12 | 37.69 |

**Table 3.** Individual and average (across one image) lesion luminance for all of the test images. The amount of *Bothrops asper* venom injected into the mice is indicated next to each batch of images.

| Luminance (cd/m<sup>2</sup>) |  |  |  |  |  |  |
| --- | --- | --- | --- | --- | --- | --- |
| Venom (µg) | Mouse 1 | Mouse 2 | Mouse 3 | Mouse 4 | Average | Std. Dev. |
| 0.5 | 0.00 | 0.00 | 0.32 | 0.39 | 0.18 | 0.18 |
| 1 | 0.24 | 0.27 | 0.38 | 0.22 | 0.27 | 0.06 |
| 2 | 0.19 | 0.27 | 0.21 | 0.23 | 0.23 | 0.03 |
| 4 | 0.07 | 0.10 | 0.09 | 0.11 | 0.09 | 0.02 |
| 8 | 0.10 | 0.09 | 0.07 | 0.07 | 0.08 | 0.01 |
| 16 | 0.10 | 0.04 | 0.09 | 0.11 | 0.09 | 0.03 |

**Table 4.** Individual and average (across one image) lesion haemorrhagic units for all of the test images. The amount of *Bothrops asper* venom injected into the mice is indicated next to each batch of images.

| Haemorrhagic units (HaU) |  |  |  |  |  |  |
| --- | --- | --- | --- | --- | --- | --- |
| Venom (µg) | Mouse 1 | Mouse 2 | Mouse 3 | Mouse 4 | Average | Std. Dev. |
| 0.5 | 0.00 | 0.00 | 19.56 | 5.93 | 6.37 | 7.99 |
| 1 | 42.01 | 49.98 | 18.09 | 40.34 | 37.60 | 11.84 |
| 2 | 74.79 | 14.43 | 79.00 | 44.38 | 53.15 | 26.04 |
| 4 | 154.67 | 90.69 | 154.71 | 63.15 | 115.81 | 40.09 |
| 8 | 180.76 | 134.09 | 239.51 | 266.37 | 205.18 | 51.41 |
| 16 | 248.77 | 644.71 | 233.08 | 280.86 | 351.85 | 169.96 |

### 3.4 Calculation of minimum hemorrhagic dose

Using linear regression (Figure X), we obtained the function *Y* = 21.83(∓ 1.18)*X* +13.73(∓ 8.91), from which we calculated the MHD on the proposed method from our earlier work (50 HaU) [21]. This resulted in a MHD of 1.66 (SD: −0.34, +0.30).

### 3.5 Tool GUI

To ensure accessibility and easy implementation of ALOHA across research, production, and quality control laboratories, a graphical user interface was developed (https://github.com/laprade117/ALOHA). Our tool can be used to quickly upload an image and receive statistics on the lesion area, luminance, and HaU for each mouse in the image (Fig. 5).

**Figure 5.**
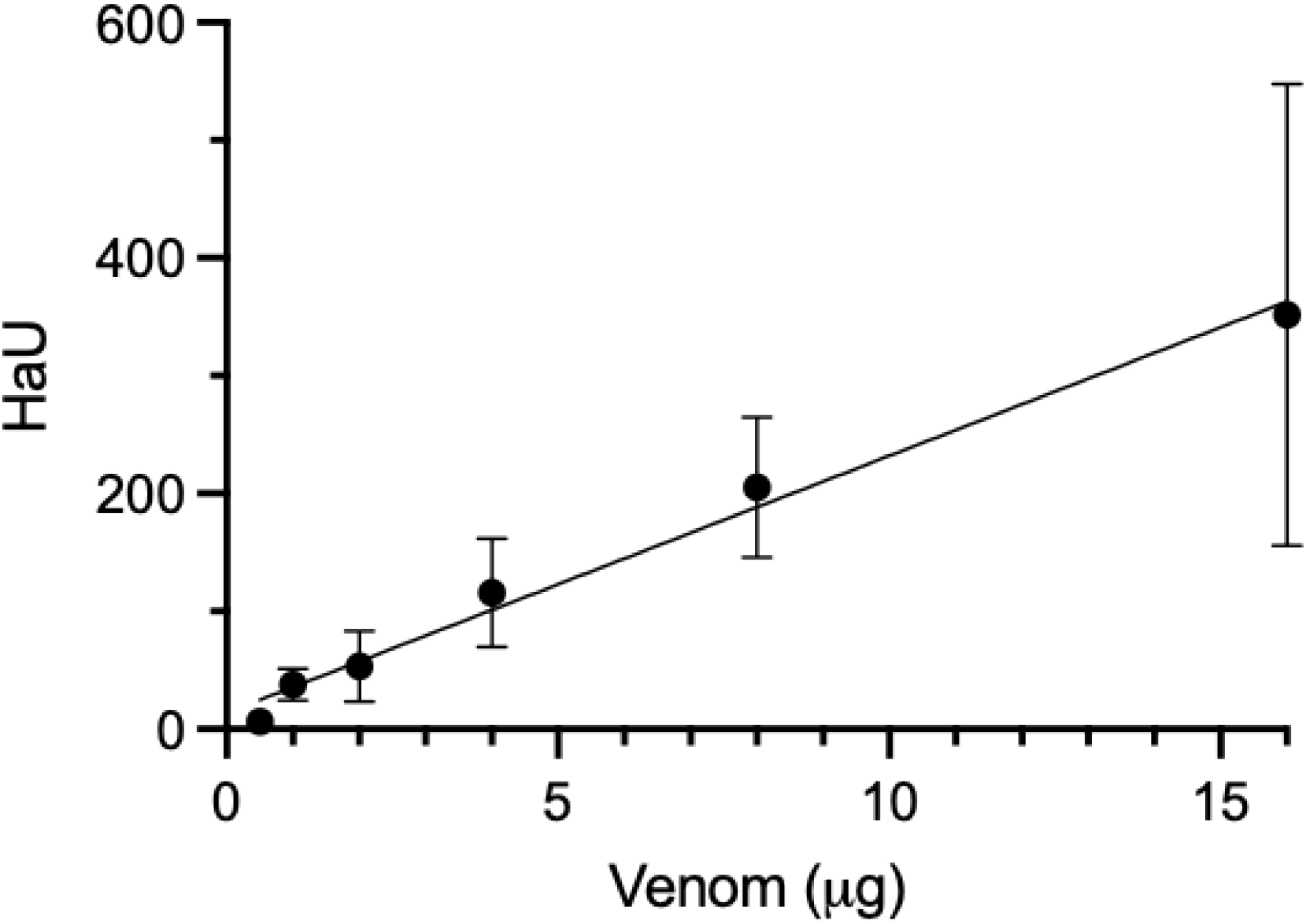
Linear regression of the means calculated from table 4, with standard deviations shown as error bars. The y-axis shows the HaUs and the x-axis shows the amount of venom used for the corresponding HaU level. Analysis was carried out using GraphPad Prism version 9.2.0.

**Figure 6.**
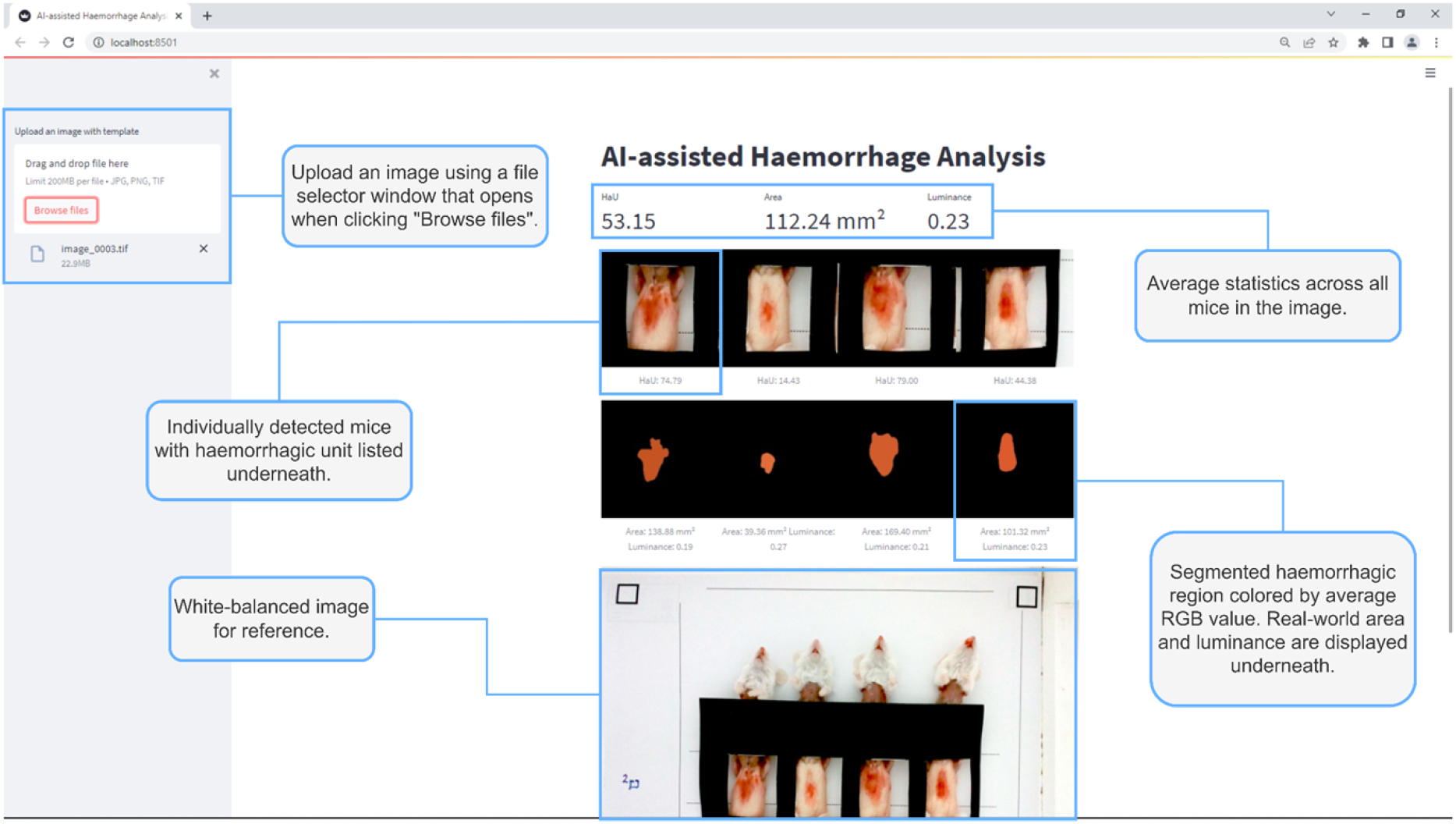
Image of ALOHA’s web interface. By uploading an image file of the experiment, the tool will, within seconds, provide the white balanced reference image, the individual lesions it detected, and how it decided to segment them, as well as all relevant data on lesion area, luminance, and HaUs.

## 4. Discussion

Haemorrhage is one of the key pathophysiological manifestations of snakebite envenomings, particularly those inflicted by species of the family Viperidae [1,11]. Therefore, the preclinical assessment of antivenom efficacy includes the evaluation of the neutralisation of haemorrhagic activity [27]. This results in both a need for robust and reliable, but also rapid assay approaches, while limiting required resources, which would allow their implementation in diverse laboratory settings. In our earlier study, we were able to improve the classical rodent skin test by increasing the accuracy of haemorrhage characterisation and reducing the time required for analysis in comparison to the original approach, as well establish that HaUs accurately reflect haemoglobin content analysis^18^. Therefore, our previous tool has been used in a series of later publications [28–37]. Nevertheless, our prior approach still remained time consuming, and required familiarisation with a new software and access to a colour pantone. Furthermore, it was subject to human error as lesions were manually annotated, which required training and would still differ from one individual to another.

In this study, we aimed to address these shortcomings by implementing a fully automated analysis pipeline, aided by vision AI, i.e. U-Net. We found that our tool, ALOHA, was able to rapidly and robustly assess the training images that covered a range of haemorrhagic lesion severities. We observed consistent white balancing, error-free scaling, and accurate segmentation. Notably, segmentation was conducted on a limited dataset to minimise animal usage prior to validation of this approach. Nevertheless, U-Net has demonstrated, across many studies [22,38–40], its robustness despite small training datasets, as was the case in our study; segmentation consistently aligned with our expert opinion, even when artificially manipulating the images to simulate a range of different laboratory/lighting conditions for a harsher image testing environment. Additionally, the HaU values calculated across the test images fall within the same range as prior findings [21,28,32,37], allowing for easy comparison and the prior validated suggestion of 50 HaU as MHD [21].

To ensure optimal accessibility and ease of implementation into existing workflows, we developed a GUI-based web tool that allows users to conduct fast and accurate analyses of haemorrhagic lesions. Using a smartphone, it takes less than a minute to take a photo, upload the image, and receive accurate information on the severity of a venom-induced haemorrhagic lesion in mice. This substantially decreases the analysis time required, from hours, to just a few minutes. Furthermore, the ease-of-use significantly boosts the accessibility of our method and provides a standard tool to be used across labs that does not require training or prior knowledge on lesion assessment.

Despite the benefits ALOHA holds, some possible limitations may exist. In properly illuminated environments, the white balancing performs accurately. However, in shadowed, or strangely lit environments, the white balancing may not perform as well. To mitigate this, it is recommended to photograph in bright, uniformly lit environments. It is especially important to avoid casting shadows on or covering the black squares on the template paper for optimal results. Furthermore, due to the translucency of a single sheet of paper, it is best to avoid placing the template on a brightly colored table or desk when photographing. Scaling is computed via information obtained from the black squares at the corners of the template paper. This computation assumes that the black squares are perfect 10 mm x 10 mm squares. Thus, for the most accurate results, it is best to use flat unwrinkled paper and photograph directly from above, so that there is the least amount of distortion applied to the black squares.

## 5. Conclusion

With ALOHA, we introduce a new algorithm for the assessment of venom-induced skin haemorrhage in mice by relying on machine learning guided image analysis approaches. This algorithm eliminates the risk of human biases in assessing lesion areas and increases the speed of analysis substantially. Furthermore, its open access web-based graphical user interface makes it easy to use and implement in laboratories across the globe. This markedly decreases the resources required for a given analysis, such as analysis and training time, and ensures reproducibility of the results.

## Acknowledgments

TP Jenkins is the grateful recipient of funding from the European Union’s Horizon 2020 research and innovation program under the Marie Sklodowska-Curie grant agreement no. 713683 (COFUNDfellowsDTU). A.H.L. acknowledges funding support from the Villum Foundation (Grant No. 00025302) and the European Research Council (ERC) under the European Union’s Horizon 2020 research and innovation programme (Grant agreement No. 850974).

## Author Contributions

All authors contributed to writing and editing the manuscript.

---

**Supplementary Figure S1:** A4 printout template to be used for haemorrhage assays.

**Supplementary Figure S2:** A4 printout cover sheet to be used for haemorrhage assays.

**Supplementary Figure S3:** Graphical illustration of how to use the ALOHA tool. First you open the Google colab tool (1.), then follow the instructions and connect to the server (2.), open the menu bar if not already visible, and select Runtime -> Restart and run all (4.). After a brief wait you use the link provided at the bottom of the page and follow the instructions.

